# A new analytical pipeline for the study of the onset of mammary gland oncogenesis based on mammary organoid transplantation and organ clearing

**DOI:** 10.1101/860270

**Authors:** Emilie Lagoutte, Clémentine Villeneuve, Vincent Fraisier, Denis Krndija, Marie-Ange Deugnier, Philippe Chavrier, Carine Rossé

## Abstract

Metastasis formation is a multi-step process starting from the dissemination of transformed carcinomatous cells from the primary tumor and could occur at a very early stage of oncogenesis, before primary tumor detection. The adult mammary gland provides a unique model to investigate epithelial cell dissemination processes. Tissue clearing techniques allow imaging samples of large volume. uDSICO clearing, one of the latest tissue clearing technique developed, provides optical imaging of whole organ due to organ clearing and tissue size reduction. We wanted to take advantage of this technique to study rare events occurring in vivo.

Here, we have established a new analytical pipeline exploiting the regenerative properties of the mammary epithelium following orthotropic transplantation of organoids together with the uDISCO organ size reduction and clearing method to study early cell dissemination in the mammary gland. As proof of concept, we analyzed the localization of epithelial cells overexpressing the oncogenic protein atypical protein kinase C iota (aPKCi^+^) in the normal mammary gland and we were able to visualize epithelial aPKCi^+^ cells, surrounded by normal epithelial cells, escaping from the normal mammary epithelium and disseminating into the surrounding stroma.

---

Metastasis formation is a multi-step process starting from the dissemination of transformed carcinomatous cells from the primary tumor (Hanahan and Weinberg, 2011; Nowell, 1976). Recently, several studies suggested that metastasis formation could occur at a very early stage of oncogenesis, before primary tumor detection (Harper et al., 2016; Hosseini et al., 2016). However, little is known about the mechanism(s) involved and regarding the behaviour and fate of transformed cells in the context of an otherwise healthy epithelium.

The adult mammary gland provides a unique model to investigate epithelial cell dissemination processes. The mammary epithelial bilayer is composed of luminal and basal myoepithelial cells, sitting on a basement membrane that separates the epithelium from the surrounding stroma (Macias and Hinck, 2012). The mammary stroma, also known as mammary fat pad, comprises multiple cellular elements, including adipocytes, fibroblasts, hematopoietic cells, lymphatic and blood vessels.

Exploring cell dissemination in the context of the whole mammary gland is challenging and remains poorly compatible with classical histological techniques based on staining of thin tissue sections or even thicker tissue slices. Here, we have established a new analytical pipeline exploiting the remarkable regenerative properties of the mammary epithelium following orthotopic transplantation of epithelial fragments or organoids (Deome et al., 1959; Jarde et al., 2016) together with the uDISCO organ clearing and size reduction method (Pan et al., 2016). Combination of these two powerful approaches enabled imaging the whole mammary gland as a mosaic of confocal z-stacks of images, providing whole organ reconstruction with a cellular-to-subcellular resolution. This allowed us to analyse the global morphology of the gland and the composition of the microenvironment while detecting epithelial or stromal localization of fluorescently tagged cells previously incorporated into the transplanted mammary organoids.

As a proof-of-concept, we applied this new pipeline to the analysis of the in vivo behaviour of mammary luminal cells overexpressing the atypical protein kinase C iota (aPKCi). aPKCi is an oncogenic protein (Fields and Regala, 2007; Parker et al., 2014; Regala et al., 2005), which is overexpressed in many carcinomas; we and others reported that aPKCi overexpression is associated bad prognosis/outcome in breast cancer (Awadelkarim et al., 2012; Rosse et al., 2014). Additionally, we recently found that overexpression of aPKCi (aPKCi^+^) in luminal epithelial cells is sufficient to trigger basally-oriented cell extrusion of aPKCi^+^ cells from the normal mammary epithelium (Villeneuve et al., 2019). Although not proven, it is possible that once they have reached the stroma, basally-extruded aPKCi^+^ cells potentially invade distant tissues.

We first adapted the uDISCO technique (Pan et al., 2016) to study the whole mammary gland samples. To improve the reduction of the size of the sample and to preserve over months antibody labelling, the uDISCO technique was preferred from older methods as 3DISCO (Erturk et al., 2012) and iDISCO (Renier et al., 2014). To this purpose, we performed an intracardiac perfusion to replace the animal’s blood by phosphate buffered saline (PBS) solution followed by an intra-cardiac fixation by injecting PBS – 4% PFA (Figure 1a – steps 1 and 2). The mammary glands #4 were sampled (Figure 1a - step 3), processed for immunostaining (Figure 1a - step 4) and then processed for size reduction and clearing (Figure 1a - step 5). Clearing and a ∼40% size/volume reduction of the whole mammary gland were observed (Figure 1b-c). We then analysed if the general organisation and the apico-basal polarity of the whole mammary epithelium were preserved by immunostaining (Figure 1c-f and Movie 1). After reduction, we were able to visualise a large portion of the mammary gland by performing mosaic reconstruction of confocal z-stacks of images (Figure d-f). The luminal epithelial cell layer (E-cadherin^+^ or keratin 8^+^) was surrounded by the myoepithelial cell layer (keratin 5^+^) outlined by the laminin 522-positive basement membrane (Figure 1d/f and Movie 1). Thus, we conclude that the organisation of the double myoepithelial and luminal epithelial cell layers was preserved as compared to immunostaining of tissue slices without uDISCO treatment (Figure 1c/d and Movie 1). The apico-basal polarity of the epithelial cells was preserved (Figure 1e) as well as the basement membrane organisation around the ducts. The epithelial cell surface and the nucleus surface area of epithelial cells were reduced by 50% (Figure 1g/h). However, the ratio of nucleus area to epithelial cell surface area was smaller after uDISCO treatment (0.51, compared to 0.6 for non-cleared tissue) suggesting that the nucleus size is slightly more reduced compared to the cytoplasm (Figure 1i). Altogether, our data show that the uDISCO clearing of the mammary gland results in a two-fold reduction in the size of the organ while preserving its organization. In addition, a high organ transparency is achieved, enabling low-background immunostaining and deep-tissue imaging by classical confocal microscopy.

**Figure 1.**
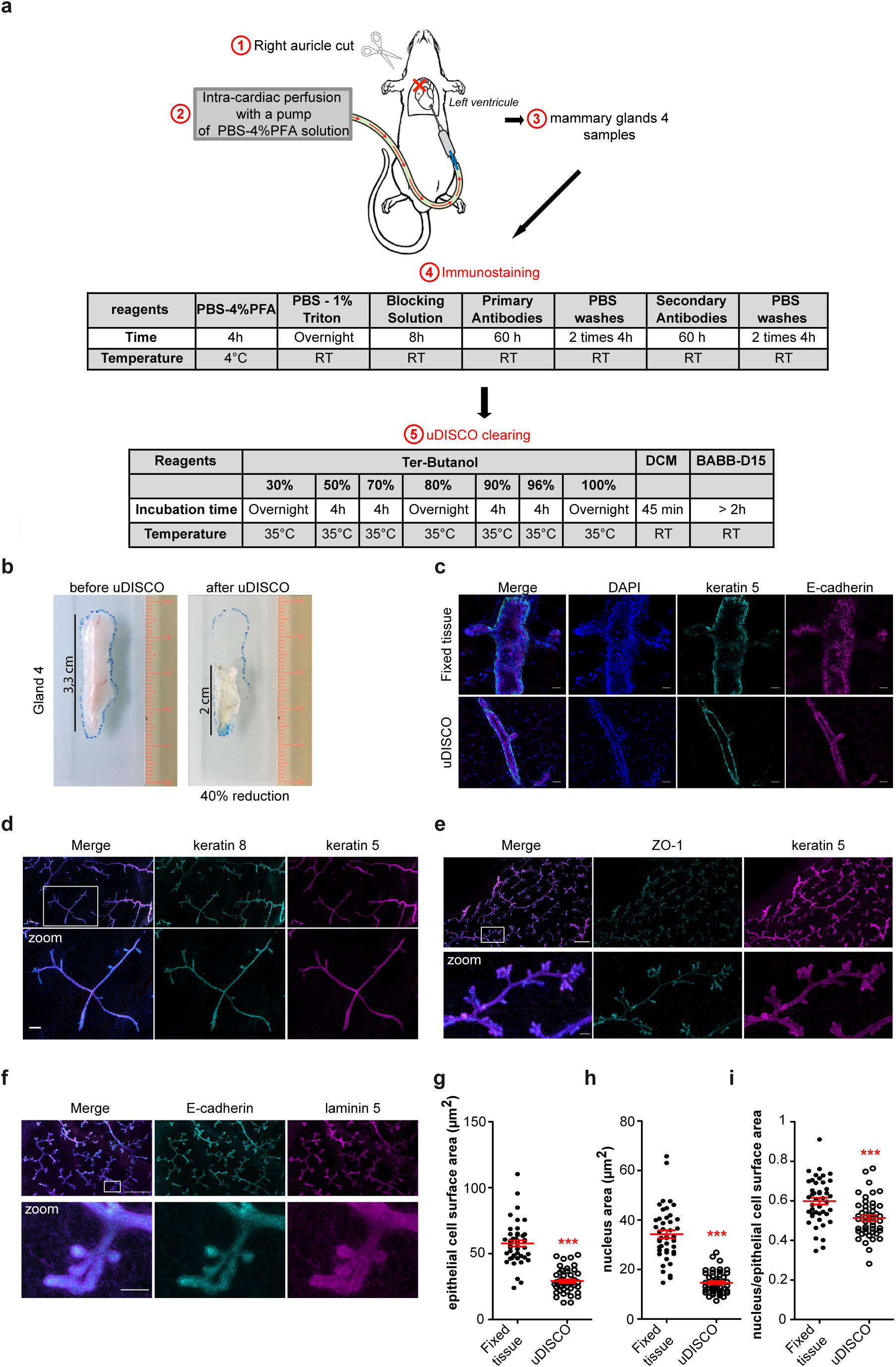
Mouse mammary gland clearing using uDISCO technique. (a) Scheme of mouse mammary gland uDISCO clearing steps. (b) Images of the mammary gland 4 from an FVB WT mouse before and after uDISCO clearing. (c) Confocal images of fixed (upper line panels) or fixed followed by uDISCO treatment mouse mammary glands 4 (lower line panels). Myoepithelial and luminal cells were respectively stained with anti-keratin-5 (cyan) and anti-E-cadherin (magenta) antibodies. Scale bars, 20µm. (d) z-projection of mosaic reconstruction of 145 planes of 2µm of confocal images of cleared mouse gland 4 (290µm thickness). Myoepithelial and luminal epithelial cells were stained with anti-keratin-5 (magenta) and anti-keratin-8 (cyan) antibodies, respectively. Scale bars, 100µm. (e) z-projection of mosaic reconstruction of 102 planes of of 3µm confocal images of cleared mouse gland 4 (306µm thickness). Myoepithelial and luminal epithelial cells were stained with anti-keratin-5 (magenta) and anti-ZO-1 (cyan) antibodies, respectively. Scale bars, 500µm (upper line panels) and 50µm (zoom panels). (f) z-projection of mosaic reconstruction of 90 planes of of 3µm confocal images of cleared mouse gland 4 (270µm thickness). Luminal epithelial cells and the basement membrane were stained with anti-E-cadherin (cyan) and anti-laminin522 (magenta) antibodies, respectively. Scale bars, 500µm (upper line panels) and 50µm. (g) Quantification of epithelial cell surface area of fixed (fixed tissue) or fixed followed by uDISCO treatment (uDISCO) mouse mammary glands 4. A Mann Whitney test was performed (***p < 0.001). (h) Quantification of epithelial nucleus surface area of fixed (fixed tissue) or fixed followed by uDISCO treatment (uDISCO) mouse mammary glands 4. A Mann Whitney test was performed (***p < 0.001). (j) Quantification of the ratio between epithelial nucleus surface area and epithelial cell surface area fixed (fixed tissue) or fixed followed by uDISCO treatment (uDISCO) mouse mammary glands 4. For each condition, 45 cells from 3 independent experiments were quantified. An unpaired t-test was performed (***p < 0.001).

Our aim was to visualise isolated rare aPKCi^+^ cells in a healthy tissue to understand the onset of oncogenesis and cell dissemination program. We focused our study on the effect of aPKCi overexpression in luminal cells. To this aim, mammary gland fragments were purified as multicellular organoids and infected with lentiviral particles at low m.o.i. to generate GFP or GFP-aPKCi-expressing cells at low frequency (Figure 2a). Using this strategy, we could overexpress aPKCi in a limited number of mammary luminal cells as shown in ((Villeneuve et al., 2019).

**Figure 2.**
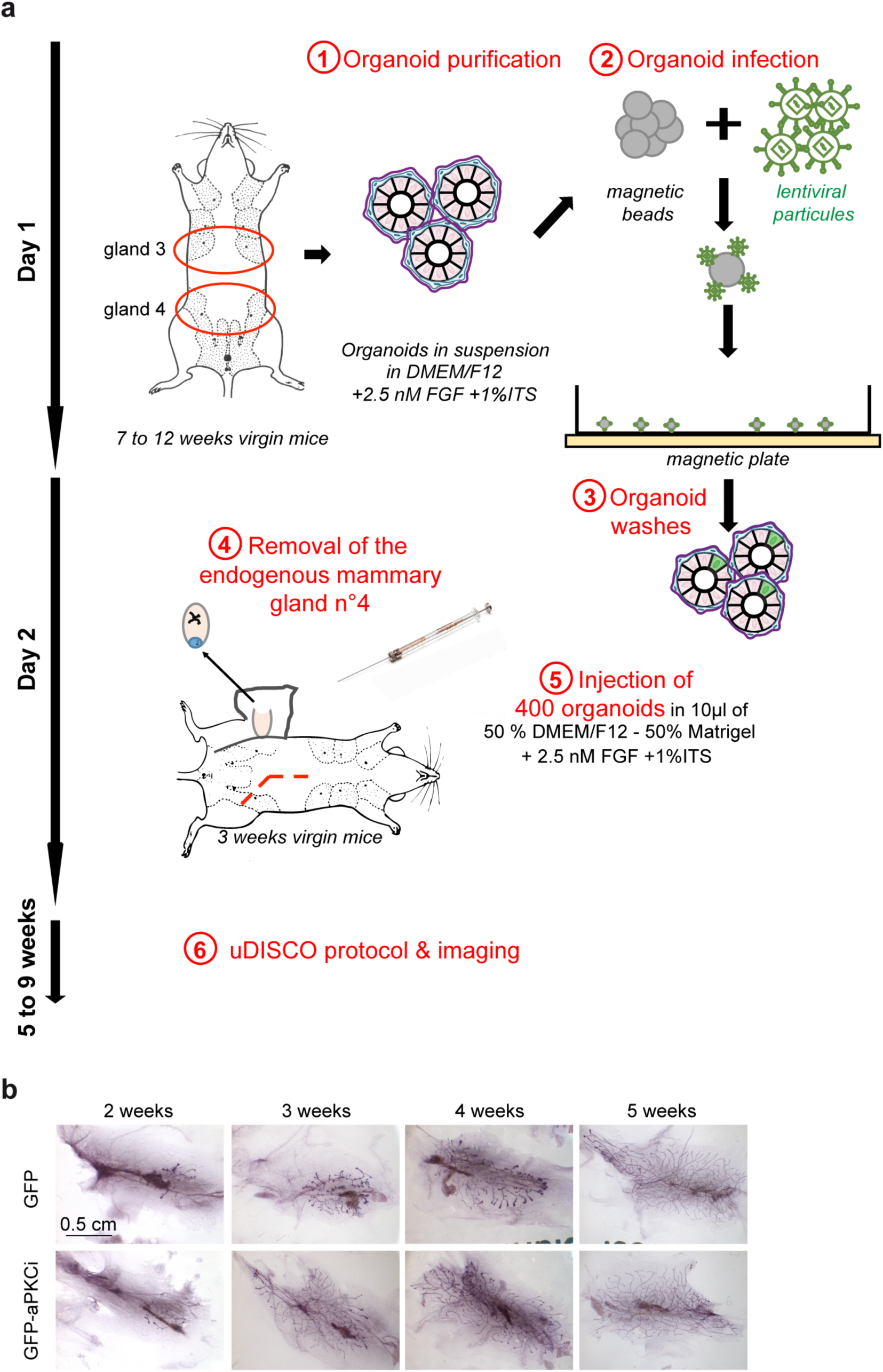
Mouse mammary infected organoids transplantation experiment. (a) Scheme of mouse mammary infected organoid transplantation steps. (b) Images of whole-mount carmine-stained regenerated mammary glands from infected GFP^+^ or GFP-aPKCi^+^ organoids sampled at different weeks post-transplantation.

Mammary organoid purification and infection were achieved within one day ex vivo. The day after, control GFP (GFP^+^) or GFP-aPKCi (GFP-aPKCi^+^) organoids were transplanted into the cleared mammary fat pad of a 3-week-old host female (Figure 2a). Both donor and host mice were of identical FVB genetic background. This in vivo grafting assay, originally described by De Ome et al. in 1959, is still widely used by the scientific community to study developmental features of the mammary gland as well as mammary tumour initiation and progression. Here, to minimize both inter-individual variability linked to the hormonal cycle and the number of animals, control GFP^+^ and mutant GFP-aPKCi^+^ organoids were co-laterally injected into the fat pad of the same recipient mouse. After five weeks, grafted organoids successfully regenerated a mammary ductal network, typical of a virgin mouse. We did not observe any visible morphological differences between the mammary epithelial trees derived control GFP^+^ and mutant GFP-aPKCi^+^ organoids (Figure 2b).

Having settled the transplantation assay from mammary organoids, we then applied the uDISCO protocol for further imaging of the regenerating epithelium. Five to ten weeks post-transplantation, intracardiac fixation of the host mice was performed as previously described (Figure 1a and material & methods sections), followed by double immunostaining for keratin 8/laminin5 (Figure 3a/b) or phospho-MLC2/E-cadherin (Figure 3c/d). Both keratin 8 and E-cadherin are specifically expressed by luminal cells, whereas phospho-MLC2 is a myoepithelial and endothelial cells marker. Laminin 5 is a major component of the basement membrane. The regenerated mammary gland was then cleared using the uDISCO procedure described above. Analysis of the GFP signal, together with the lineage specific markers, showed that mammary ducts contained only a few GFP^+^ or GFP-aPKCi^+^ cells surrounded by Wild-Type (WT) luminal cells (Figure 3 and movies 1-3).

**Figure 3.**
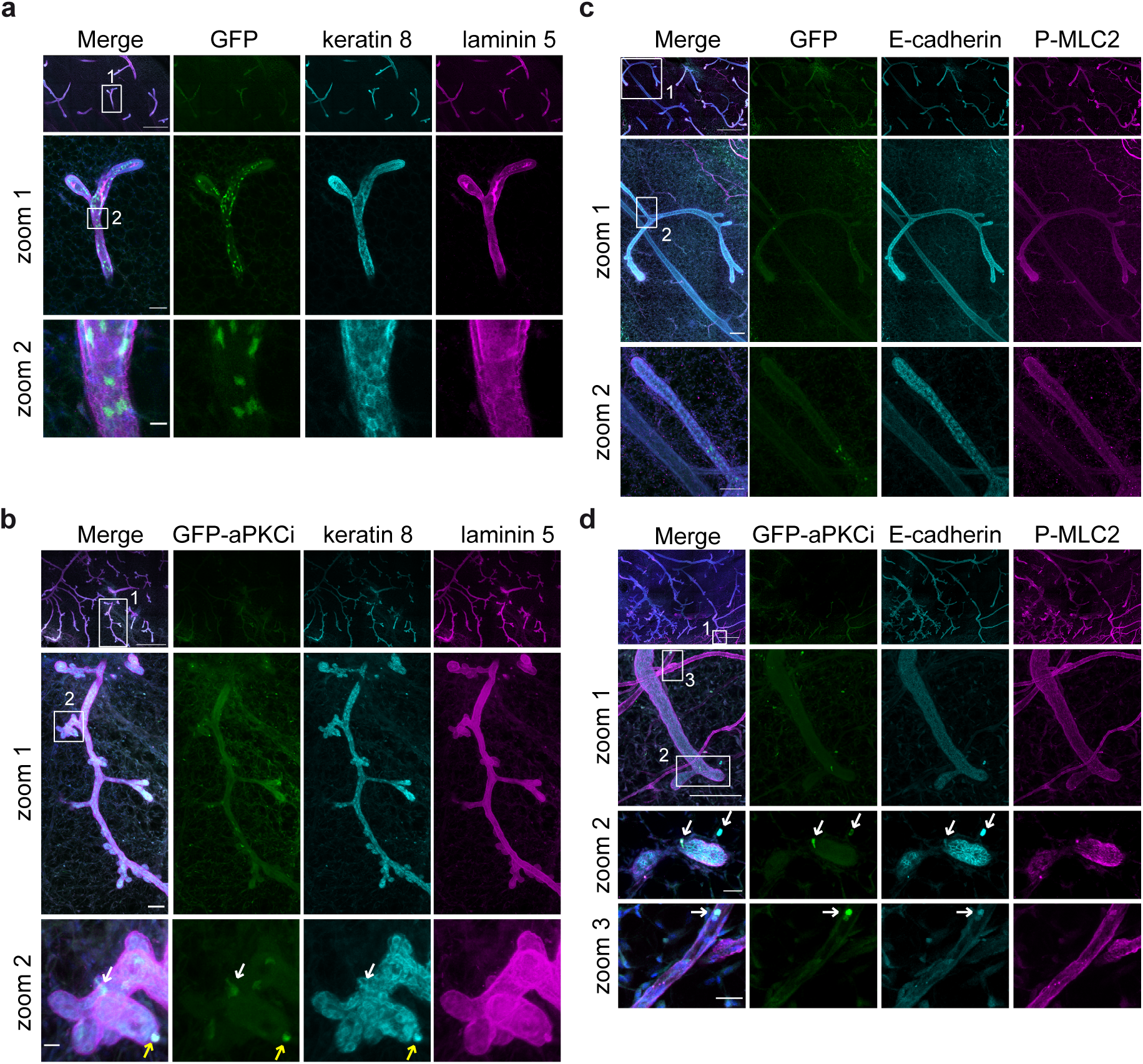
Combination of organ clearing and mouse mammary organoid transplantation allows the detection of aPKCi^+^ luminal epithelial cells breaching the basement membrane and disseminating into the surrounding stroma. (a-b) Confocal images of regenerated mammary glands from GFP^+^ (a) or GFP-aPKCi^+^ (b) infected mouse organoids. The basement membrane and epithelial cells were stained with anti-laminin 5 (magenta) and anti-keratin 8 (cyan) antibodies, respectively. Scale bars, 500µm (upper line panels), 50µm (zoom 1) and 10µm (zoom 2). (a) Upper line panels: z-projection of mosaic reconstruction of 93 confocal planes (186µm thickness); zoom 1 and 2: z-projection of 15 confocal planes (30µm thickness). (b) z-projection of mosaic reconstruction of 183 confocal planes (366µm thickness). The white arrows indicate GFP-aPKCi^+^ luminal cells breaching the basement membrane. The yellow arrows indicate GFP-aPKCi^+^ luminal epithelial cells completely extruded from the mammary ducts. (c-d) Mosaic reconstruction of confocal images of ten weeks post transplantation regenerated mammary glands from GFP^+^ (c) or GFP-aPKCi^+^ (d) infected mouse organoids. Samples were stained with anti-P-MLC2 (magenta) and anti-E-cadherin (cyan) antibodies. The P-MLC2 staining allows the detection of the myoepithelial cell layer and the blood vessel. The E-cadherin staining allows the detection of the mammary luminal cells. (c) z-projection of 152 confocal planes (304µm thickness). Scale bars, 500 µm (upper line panels), 100µm (zoom1) and 50µm (zoom2). (d) Upper line panels and zoom1: z-projection of 176 confocal planes of confocal images (352µm thickness). Zoom2: z-projection of 24 confocal planes (48µm thickness) showing GFP-aPKCi^+^ luminal epithelial cells escaping from the mammary ducts and in the stroma. Zoom3: z-projection of 18 confocal planes (36µm thickness) showing GFP-aPKCi^+^ luminal epithelial cells in blood vessels. Scale bars, 500µm (upper line panels) and 100µm (zoom1) 20µm (zoom2&3).

As for the normal mammary gland, a significant reduction of total cell surface area (about 45%) and nucleus surface area (about 35%) (Sup. Fig1a/b) was observed without a significant reduction of nucleus/total cell surface area ratio (Sup. Fig1c). All GFP^+^ epithelial cells were localised into the ducts (Figure 3a), whereas GFP-aPKCi^+^ cells, positive for keratin 8, were able to breach the basement membrane (Figure 3b - Movie 2). These data, in line with our previous study (Villeneuve et al., 2019), show that GFP-aPKCi^+^ cells could leave the duct by transmigration through the basement membrane. The phospho-MLC2 staining, in combination with E-cadherin staining, allowed us to detect both the mammary ducts and the blood vessels (Figure 3c/d). Again, all GFP^+^ epithelial cells were localized into the ducts (Figure 3c - Movie 3), whereas we could detect some GFP-aPKCi^+^ cells, positive for E-Cadherin, able to breach the myoepithelial cell layer (labelled by an anti-phospho-MLC2) surrounding the luminal epithelial E-cadherin^+^ cells. Moreover, some of the GFP-aPKCi^+^ cells were found disseminated into blood vessels (also labelled by an anti-phospho-MLC2; Figure 3d and Movie 4). Notably, the “escaping” GFP-aPKCi^+^ cells were positive for the luminal markers, keratin 8 and E-Cadherin, suggesting that they did not undergo epithelial-mesenchymal transition (EMT) upon migration.

Here, we have shown a novel imaging-based method to study the onset of mammary gland oncogenesis, allowing whole-organ imaging analysis with (sub)cellular resolution while preserving a physiological context. Through combined use of infected organoids transplantation and uDISCO clearing we were able to detect and analyse rare cellular events occurring *in vivo*. Using this approach, we showed that aPKCi overexpression promotes basal extrusion of mammary luminal epithelial cells and cell invasion into the stroma, pointing out to a potential path for cell dissemination without/before formation of a primary tumour. However, while this technique is compatible with immunolabelling allowing detection of endogenous proteins as well as tagged proteins, some autofluorescence in the GFP channel could disturb the visualisation of weak signals. In this line, it might be useful to test this adapted protocol made for the brain (Li et al., 2018) in our model to reduce this issue. Summary of advantages and limitations of our method are listed in Table 1.

**Table 1.** Advantages and limitations of the method.

| **Advantages** |
| --- |
| - A high quality mosaic image<br>- 3D view of the whole organ with preservation of the entire structure and the intracellular organization<br>- Detection of rare events in the whole mammary gland<br>- Identification of various components of the mammary gland, such as epithelial and myoepithelial cells, the basement membrane and blood vessels<br>- Intact immune system |
| **Limitations** |
| - Autofluorescence of the mammary gland after clarification in the GFP channel<br>- Long processing time of the samples. Clearing and immunostaining processes take around 12 days as well and the confocal acquisition takes 6h to 12h<br>- Requirement for powerful computers due to large size of the images (25 to 100 Go).<br>- Lymphatic cell dissemination can not be studied due to the lymph node removal |

The uDISCO clearing has been previously described in other organs (Pan et al., 2016) and it could be interesting in our study to perform this technique on potential colonization sites as the lung and the brain. Moreover, this combination of techniques could be used to study the ability of mammary genetically-modified stem cells to regenerate the mammary gland. This method is adapted to analyse the impact of the stroma on the mammary gland development or on tumor progression using genetically-modified mammary organoids.

## Material & Methods

### Isolation of the mammary epithelium and generation of organoids

The mammary epithelium isolation protocol is similar than the method described in (Ewald, 2013; Nguyen-Ngoc et al., 2012). Quickly, the two glands 3 and 4 of both sides of the mice were collected and lymphatic ganglions from glands 4 were removed. Then, the mammary glands were cut with a scalpel about 50 times and the small pieces were transferred into 25 mL of a collagenase solution (Table S1) (volume up to 2 mice). The collagenase solution containing the glands was incubated for 1h at 37 ° C with shaking at 120 rpm and a manual shake at 30 minutes.

After this step, all the tubes and tips were coated with PBS - 0,3% BSA. The collagenase solution containing mammary gland fragments was transferred into a glass tube (tube #1) and centrifuged at 500g for 10 minutes at 25°C. Three layers were obtained in the tube: a fatty layer at the top, an aqueous layer in the middle and the basal layer containing the epithelial material at the bottom. The fatty layer was transferred into a second tube (tube #2) of 25 mL of a collagenase solution and was incubated 20 minutes at 37 °C at 120 rpm.

Meanwhile, the tube #1 was centrifuged at 500g for 10 minutes at 25°C; the pellet was resuspended in 10 mL of organoid medium without ITS (Table S1).

At the end of the 20 minutes, the tube #2 was centrifuged in a glass tube for 10 minutes at 25°C at 500g. The pellet was resuspended in 2 mL of organoid medium without ITS before being transferred into the tube #1. The tube #1 was centrifuged at 500g for 10 minutes at 25°C. The pellet was resuspended in 4mL of DNAse solution (Table S1) and the tube was mixed by hand during 2 to 5 minutes at room temperature (RT). Then, 6 mL of organoid medium without ITS was added. The tube was centrifuged at 500g during 10 minutes at 25°C and the purification of the pellet of organoids was performed, consisting in 2 round of 2 minutes centrifugation at 500g for 10 minutes followed by 3 pulses of centrifugations at 500g in 10 mL of organoid medium.

The last pellet was resuspended in the appropriate volume of organoid medium supplemented with 1% ITS and 2.5 nM FGF (about 600-800 organoids/ml). 700 µL / well was distributed in non-adherent 24 wells plate.

### Low-efficiency lentiviral infection of organoids

#### Production of lentiviral particle

##### One day before transfection

The HEK 293T cells were grown in 25cm2 culture flasks on appropriate medium (Table S1) (2 flasks per condition) to obtain 60-70% confluence on the day of transfection at 37ºC and 5% CO2.

##### Day of transfection

The medium was changed 2h before transfection. A mix of 400 µl of OptiMem medium and 18 µl of GeneJuice was made and mixed by vortex (Tube A). The mix was incubated at room temperature for 5 minutes. In parallel, 0.9 µg pCMV VsVg, 2.1 µg of psPAX 2 and 3 µg of vector containing the protein to express were gently mixed by up & down with 400 µl of OptiMem medium (Tube B).

The tubes A and B were mixed up & down in a total volume of 800µl. The final mix was incubated at room temperature for 20 minutes. 400µl of the mix was added drop by drop per flask. Cells were incubated during 60 hours at 37ºC and 5% CO2.

#### Concentration of lentiviral particles

The supernatant containing the lentiviral particles was sampled and filtered with a 0.45µm syringe filter. After filtration, the supernatant was concentrated with Lenti-X concentrator (1 volume of Lenti-X concentrator for 3 volumes of supernatant) and the mix was incubated 2h at 4°C. Then, the mix was centrifuged during 45 min at 4°C at 1500g. The pellet was resuspended in organoid medium to concentrate the viral particles solution 50 times. The solution was aliquoted and frozen at −80°C until the experiment was performed. Cycles of frozen & thaw should be avoided.

#### Infection of the organoids

Magnetic beads were added to the viral particle solution (Table S2) (1/10eme of the volume of virus particle solution) and incubated 20-25 minutes at RT. Then the mix magnetic bead – virus particle solution was added on organoids plated on non-adherent 24 wells plate (55µL of the mix for one well). The non-adherent 24 wells plate was put on a magnet for 1 hour 30 minutes at 37°C, 5% CO2. Then, the magnet was removed and the 24 wells plate was incubated at 37°C, 5% CO2 for 24h.

Note 1: During manipulation, all plastics were coated with PBS-0.3% BSA.

#### Preparation of the organoids for the transplantation

Organoids were transferred into a 50 mL falcon and washed three times in 5 mL of PBS - 0.3% BSA with three centrifugation cycles of 10 min at 500g at RT. Organoids were resuspended at a concentration of 400 organoids/10 µL in transplantation medium (TableS1).

### Transplantation

Between 4 and 6 animals per experiment were transplanted in order to obtain significant results per lot. The entire transplant procedure was performed under general anaesthesia. The anaesthetic solution used was Ketamine (10mg/mL) (also analgesic) - Xylazine (1mg/mL), - Flunitrazepam (1mg/mL) (benzodiazepine, which ensures a loss of consciousness and a peaceful awakening) (Table S2). This solution was diluted by two in PBS. 100µL of this solution was intraperitoneally injected per 10gr of mouse. The anesthetized mice were placed on a warm cushion during all the operation to maintain their body temperature at 37°C. Before the operation, 200µL of physiological serum was sub-cuteanously injected to hydrate the mice. During surgery, the legs were held by a flexible adhesive that does not impede blood circulation and mice were depilated on the operated area. The skin of the abdomen was incised (an intact peritoneum was preserved) on 2 cm. The vessels were cauterized with a cauterizer and the original gland was eliminated. Then 10 µL of cells (equivalent to about 400 organoids resuspended in transplantation medium (Table S1) was injected using an ultrafine Hamilton syringe (25 µL Hamilton 702 RN). After surgery, the skin was sew using resorbable surgical threads to avoid discomfort and infections. Postoperatively, the anesthetized mice were placed on a hot plate to avoid hypothermia. Upon waking, mice were placed in a new cage with water and food on demand. During the first two weeks of storing, hydrogel cups with ground food were placed in cages to facilitate food access for mice. The mice were monitored every day for the first 4 days after transplantation, then once a week. The skin was quickly (one week approximately) healed and mice were not embarrassed in its movements.

### 3D tissue immunofluorescence

The mice were then sacrificed at the latest 5 to 10 weeks post-transplantation and the mammary glands 4 were collected. For mammary gland immunofluorescence, without clearing and reduction, tissue immunofluorescence was performed as described in (Villeneuve et al., 2019).

Note 2: The fixed and immunostained tissues could not be kept more than 2 weeks. Freezing the sample would impact the structure of the gland and immunostaining.

### Histological mouse regenerated mammary gland tissue

Whole-mount carmine was performed as described (Teuliere et al., 2005).

### Intra-cardiac fixation

The animals were deeply anesthetized with Ketamine (Table S2, 80-100 mg / mL) mixed with the analgesic Xylasin (Table S2, 5-10 mg/mL) in PBS. The mixture was intra-peritoneally injected at a concentration of 50µL/10g of mouse weight. The efficiency of the anaesthesia was checked by pinching the mice, and the procedure was started if no muscle response was detected. An incision was made from the lower abdomen to the top of the thorax and the rib cage was opened. The needle of the peristaltic pump was inserted in the left ventricle of the heart and the vein returning the blood to the right atrium was severed. A solution of phosphate buffered saline (PBS) was perfused, followed by PBS - 4% paraformaldehyde (30 mL per mice).

### Staining uDISCO

#### Whole Gland Staining

The whole gland was fixed for 3-4h in PBS - 4% PFA at 4°C. The staining protocol was adapted from (Krndija et al., 2019). Briefly, the gland was permeabilised using PBS - 1% Triton-X100 overnight at RT under shaking. Directly after permeabilization, the gland was incubated in blocking solution for 8 hours in PBS - 0.2% Triton - 1% BSA - 3% FBS. Then the gland was incubated in 300µL to 500µL of the primary antibody diluted at the adequate concentration (Table S3) in PBS - 0.2% Triton at RT for at least 2 days. The gland was washed 3 times during the day with PBS - 0.2% Triton. The gland was then incubated with the secondary antibody diluted in PBS - 0.2% Triton at RT for at least 2 days (Table S3 for the dilution). The gland was washed 3 times during the day with PBS - 0.2% Triton.

#### Tissue clearing using the uDISCO technique

The samples were washed in 0.1M PBS for 5 minutes before clearing. All incubation steps were performed in a chemical hood with gentle rotations or shaking. The samples were covered with aluminium foil to protect them from the light. The fixed samples were incubated at 35°C overnight in tert-butanol diluted in distilled water at 30 Vol%, 4 hours in 50 Vol%, 4 hours in 70 Vol%, overnight in 80 Vol%, 4 hours in 90 Vol%, 4 hours in 96 Vol%, 4 hours in 100% tert-butanol, and then in dichloromethane for 45-60 minutes at room temperature. Eventually, the samples were incubated in BABB-D15 composed of BABB (benzyl alcohol + benzyl benzoate at a ratio 1:2) + diphenyl ether (ratio 1:15 of BABB) + 0.4% vitamin? for 2 to 6 hours until the samples became optically transparent and kept in this solution.

Note 3: If samples are not immediately cleared after immunostaining, they should be stored at 4°C in PBS - 0.2% Triton.

### Imaging

Confocal images of cleared samples or mammary tissues were performed on a CLSM Leica TCS SP8 inverted microscope equipped with hybrid detectors HyD. Depending on the magnification, several objectives were used (20x PL ApoDry 0.75NA, 40x PL Apo oil 1.25NA objectives). Mosaic images were reconstructed using the Leica SP8 software. Images were processed using ImageJ software.

### Statistical analysis

All error bars represent the standard error of the mean (SEM). The statistical test used for each graph is mentioned in the legend of the graph. The statistical significance was defined as ***p < 0.001, ns for nonsignificant. All calculations and plots were performed and generated using GraphPad Prism software.

## Acknowledgments

The authors greatly acknowledge the Nikon Imaging Center at Institut Curie-CNRS, the PICT-IBiSA (member of the France-BioImaging national research infrastructure) for fluorescence microscopy equipment and for help with image acquisition. We are particularly grateful to the personnel of the Animal facility from Institut Curie. Members of P. Chavrier’s laboratory are thanked for helpful discussions. CV is funded by a grant from by PSL and Fondation pour la Recherche Médicale. Funding for this work was provided by grants to CR grants from *Projet Fondation ARC pour la Recherche contre le Cancer* (PJA20141201972), Canceropôle Ile de France (EMERG-1, INVADOID 2016) and GEFLUC, LABEX CellPhyBio ANR-11-LABX-0038 and by core funding from *Institut Curie* and *Centre National pour la Recherche Scientifique* to PC.

## Ethical Approval

Animal care and use for this study were performed in accordance with the recommendations of the European Community (2010/63/UE) for the care and use of laboratory animals. Experimental procedures were specifically approved by the ethics committee of the Institut Curie CEEA-IC #118 (CEEA-IC 2017-013) in compliance with international guidelines.

## References

Awadelkarim, K.D., C. Callens, C. Rosse, A. Susini, S. Vacher, E. Rouleau, R. Lidereau, and I Bieche. 2012. Quantification of PKC family genes in sporadic breast cancer by qRT-PCR: evidence that PKCiota/lambda overexpression is an independent prognostic factor. Int J Cancer. 131:2852–2862.

Deome, K.B., L.J. Faulkin, Jr., H.A. Bern, and P.B. Blair. 1959. Development of mammary tumors from hyperplastic alveolar nodules transplanted into gland-free mammary fat pads of female C3H mice. Cancer Res. 19:515–520.

Erturk, A., K. Becker, N. Jahrling, C.P. Mauch, C.D. Hojer, J.G. Egen, F. Hellal, F. Bradke, M. Sheng, and H.U. Dodt. 2012. Three-dimensional imaging of solvent-cleared organs using 3DISCO. Nat Protoc. 7:1983–1995.

Ewald, A.J. 2013. Isolation of mouse mammary organoids for long-term time-lapse imaging. Cold Spring Harb Protoc. 2013:130–133.

Fields, A.P., and R.P. Regala. 2007. Protein kinase C iota: human oncogene, prognostic marker and therapeutic target. Pharmacol Res. 55:487–497.

Hanahan, D., and R.A. Weinberg. 2011. Hallmarks of cancer: the next generation. Cell. 144:646–674.

Harper, K.L., M.S. Sosa, D. Entenberg, H. Hosseini, J.F. Cheung, R. Nobre, A. Avivar-Valderas, C. Nagi, N. Girnius, R.J. Davis, E.F. Farias, J. Condeelis, C.A. Klein, and J.A. Aguirre-Ghiso. 2016. Mechanism of early dissemination and metastasis in Her2+ mammary cancer. Nature.

Hosseini, H., M.M. Obradovic, M. Hoffmann, K.L. Harper, M.S. Sosa, M. Werner-Klein, L.K. Nanduri, C. Werno, C. Ehrl, M. Maneck, N. Patwary, G. Haunschild, M. Guzvic, C. Reimelt, M. Grauvogl, N. Eichner, F. Weber, A.D. Hartkopf, F.A. Taran, S.Y. Brucker, T. Fehm, B. Rack, S. Buchholz, R. Spang, G. Meister, J.A. Aguirre-Ghiso, and C.A. Klein. 2016. Early dissemination seeds metastasis in breast cancer. Nature.

Jarde, T., B. Lloyd-Lewis, M. Thomas, H. Kendrick, L. Melchor, L. Bougaret, P.D. Watson, K. Ewan, M.J. Smalley, and T.C. Dale. 2016. Wnt and Neuregulin1/ErbB signalling extends 3D culture of hormone responsive mammary organoids. Nat Commun. 7:13207.

Krndija, D., F. El Marjou, B. Guirao, S. Richon, O. Leroy, Y. Bellaiche, E. Hannezo, and D. Matic Vignjevic. 2019. Active cell migration is critical for steady-state epithelial turnover in the gut. Science. 365:705–710.

Li, Y., J. Xu, P. Wan, T. Yu, and D. Zhu. 2018. Optimization of GFP Fluorescence Preservation by a Modified uDISCO Clearing Protocol. Front Neuroanat. 12:67.

Macias, H., and L. Hinck. 2012. Mammary gland development. Wiley Interdiscip Rev Dev Biol. 1:533–557.

Nguyen-Ngoc, K.V., K.J. Cheung, A. Brenot, E.R. Shamir, R.S. Gray, W.C. Hines, P. Yaswen, Z. Werb, and A.J. Ewald. 2012. ECM microenvironment regulates collective migration and local dissemination in normal and malignant mammary epithelium. Proc Natl Acad Sci U S A. 109:E2595–2604.

Nowell, P.C. 1976. The clonal evolution of tumor cell populations. Science. 194:23–28.

Pan, C., R. Cai, F.P. Quacquarelli, A. Ghasemigharagoz, A. Lourbopoulos, P. Matryba, N. Plesnila, M. Dichgans, F. Hellal, and A. Erturk. 2016. Shrinkage-mediated imaging of entire organs and organisms using uDISCO. Nat Methods. 13:859–867.

Parker, P.J., V. Justilien, P. Riou, M. Linch, and A.P. Fields. 2014. Atypical protein kinase Ciota as a human oncogene and therapeutic target. Biochem Pharmacol. 88:1–11.

Regala, R.P., C. Weems, L. Jamieson, A. Khoor, E.S. Edell, C.M. Lohse, and A.P. Fields. 2005. Atypical protein kinase C iota is an oncogene in human non-small cell lung cancer. Cancer Res. 65:8905–8911.

Renier, N., Z. Wu, D.J. Simon, J. Yang, P. Ariel, and M. Tessier-Lavigne. 2014. iDISCO: a simple, rapid method to immunolabel large tissue samples for volume imaging. Cell. 159:896–910.

Rosse, C., C. Lodillinsky, L. Fuhrmann, M. Nourieh, P. Monteiro, M. Irondelle, E. Lagoutte, S. Vacher, F. Waharte, P. Paul-Gilloteaux, M. Romao, L. Sengmanivong, M. Linch, J. van Lint, G. Raposo, A. Vincent-Salomon, I. Bieche, P.J. Parker, and P. Chavrier. 2014. Control of MT1-MMP transport by atypical PKC during breast-cancer progression. Proc Natl Acad Sci U S A. 111:E1872–1879.

Teuliere, J., M.M. Faraldo, M.A. Deugnier, M. Shtutman, A. Ben-Ze’ev, J.P. Thiery, and M.A. Glukhova. 2005. Targeted activation of beta-catenin signaling in basal mammary epithelial cells affects mammary development and leads to hyperplasia. Development. 132:267–277.

Villeneuve, C., E. Lagoutte, L. de Plater, S. Mathieu, J.B. Manneville, J.L. Maitre, P. Chavrier, and C. Rosse. 2019. aPKCi triggers basal extrusion of luminal mammary epithelial cells by tuning contractility and vinculin localization at cell junctions. Proc Natl Acad Sci U S A.

