## Supplemental data for "A new analytical pipeline for the study of the onset of mammary gland oncogenesis based on mammary organoid transplantation and organ clearing"

* co-last author

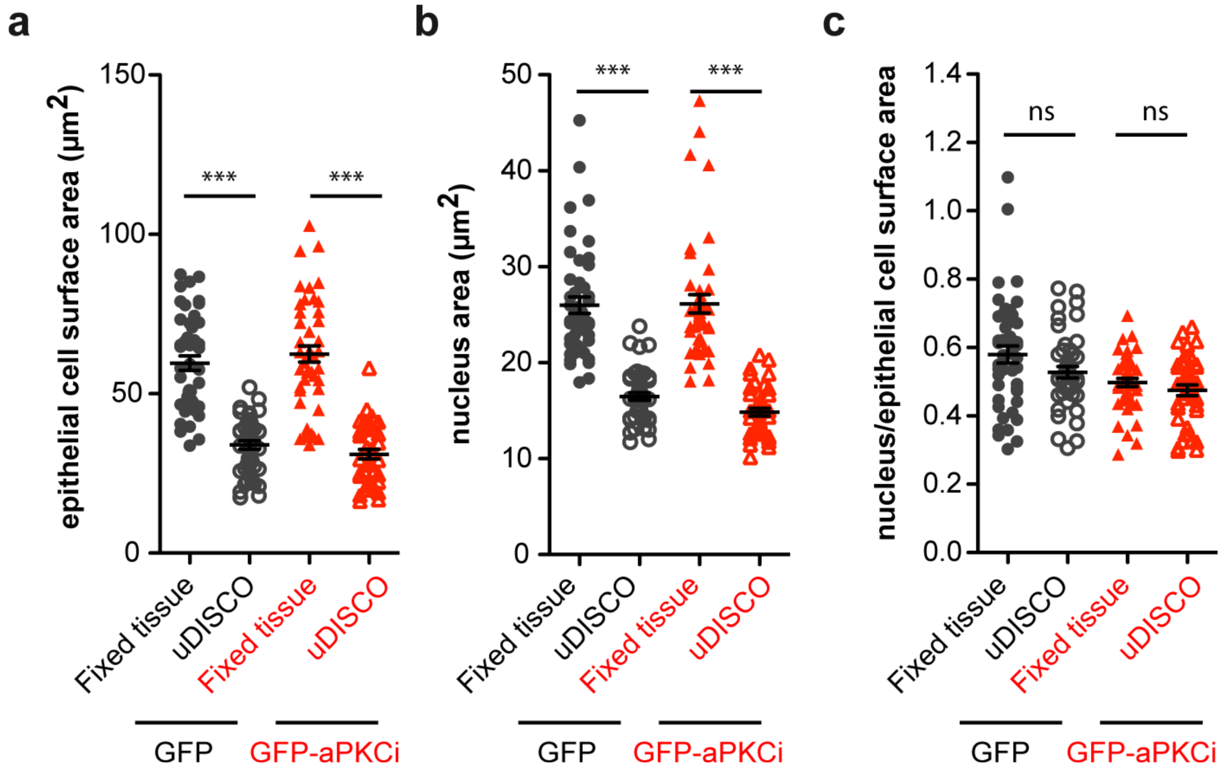


**Sup. Fig. 1. Quantification of epithelial cell surface and nucleus area of fixed tissue (fixed tissue) or fixed tissue followed by uDISCO treatment (uDISCO) from regenerated mammary glands from GFP$^+$ or GFP-aPKCi$^+$ organoids.** (a) Quantification of epithelial cell surface area. A Kruskal-Wallis test was performed (***$p < 0.001$). (b) Quantification of epithelial nucleus surface area, a Kruskal-Wallis test was performed (***$p < 0.001$). (c) Quantification of the ratio between epithelial nucleus surface area and the epithelial cell surface area. A Kruskal-Wallis test was performed (ns, nonsignificant). (a-c) For "GFP-fixed tissue", "GFP-uDISCO", "GFP-aPKCi-fixed tissue" conditions, 45 cells from 3 independent experiments were quantified. For "GFP-aPKCi-uDISCO" condition, 41 cells from 3 independent experiments were quantified.

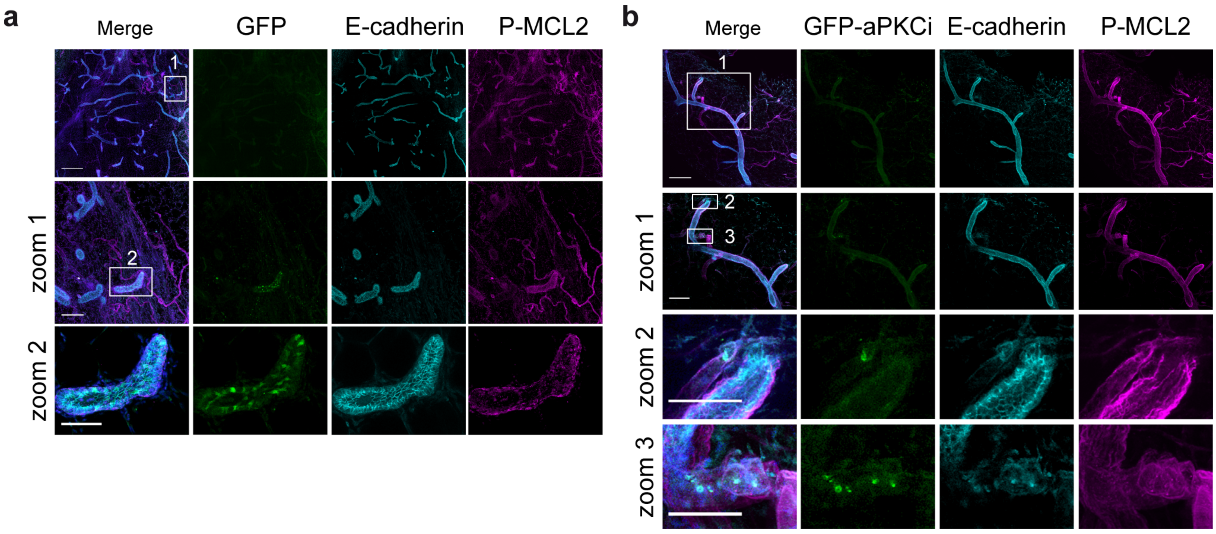


**Sup. Fig. 2. Combination of organ clearing and mouse mammary organoid - transplantation allows the detection of aPKCi$^+$ luminal epithelial cells disseminating into the mammary stroma.** Six weeks post transplantation regenerated mammary glands were stained with anti-P-MLC2 (magenta) and anti-E-cadherin (cyan) antibodies. The P-MLC2 staining allows the detection of the myoepithelial cell layer and the blood vessel. The E-cadherin staining allows the detection of the mammary luminal cells. (a) Upper line panels: z-projection of mosaic reconstruction of 102 planes of confocal images (204µm thickness) of regenerated mammary glands from GFP$^+$ infected mouse organoids. Scale bar, 500 µm. Zoom1: z-projection of 70 planes of confocal images (140µm thickness). Scale bar, 100µm. Zoom2: z-projection of 7 planes of confocal images (14µm thickness). Scale bar, 50µm. (b) Upper line panels and zoom1: z-projection of mosaic reconstruction of 97 planes of confocal images (194µm thickness) of regenerated mammary glands from GFP-aPKCi$^+$ infected mouse organoids. Scale bar, 500µm. Zoom1: z-projection of 37 planes of confocal images (74µm thickness). Scale bar, 100µm. Zoom 2&3: z-projection of 10 planes of confocal images (20µm thickness) of regenerated mammary glands from GFP-aPKCi$^+$ infected mouse organoids showing GFP-aPKCi$^+$ luminal epithelial cells in blood vessels. Scale bars, 20µm.

**Movies**

**Movie 1. 3D reconstruction of FVB mouse mammary gland 4** performed on Imaris software. The movie corresponds to the z-stack shown in Fig. 1d of 145 planes of cleared mouse gland 4 (290µm thickness) showing the localization of laminin5 (in magenta) and keratin8 (in cyan). Z-stack images were captured at 2µm interval.

**Movie 2. Detection of GFP-aPKCi$^{+}$ luminal epithelial cells breaching the basement membrane and disseminating into the surrounding stroma.** The movie corresponds to the z-stack shown in Fig. 3b - zoom2 showing the localization of laminin5 (in magenta) and keratin8 (in cyan). Z-stack images were captured at 2µm interval.

**Movie 3. Detection of GFP$^{+}$ luminal epithelial cells of regenerated mammary glands from GFP$^{+}$ infected mouse organoids:** 3D reconstruction performed on Imaris software corresponding to the figure 3c of 152 planes of confocal images (304µm thickness). Z-stack images were captured at 2µm interval.

**Movie 4. Detection of GFP-aPKCi$^{+}$ luminal epithelial cells of regenerated mammary glands from GFP-aPKCi$^{+}$ infected mouse organoids:** 3D reconstruction performed on Imaris software corresponding to the figure 3d of 176 planes of confocal images (352µm thickness). Z-stack images were captured at 2µm interval.
